# TooTranslator: Zero-Shot Classification of Specific Substrates for Transport Proteins by Language Embedding Alignment of Proteins and Chemicals

**DOI:** 10.1101/2025.11.13.688294

**Authors:** Sima Ataei, Gregory Butler

## Abstract

Transmembrane transport proteins mediate selective movement of ions and metabolites across membranes. Experimental characterization of their substrate specificity is limited. For novel class discovery with limited data, zero-shot learning aims to assign labels to test samples whose label has not been seen previously during training of the model. We introduce TooTranslator, a regression-based model that aligns embeddings from pre-trained protein (ProtBERT), chemical (ChemBERTa), and text (SciBERT) language models into a shared latent space, enabling substrate prediction by minimizing distances between protein and substrate embeddings. TooTranslator tackles zero-shot learning for the task of predicting the specific substrate of transmembrane transport proteins. Models using four loss functions are evaluated on protein test sets with seen and unseen substrates. We find no statistically significant difference in performance of the four loss functions. For tests with seen substrates, the models compare with the state-of-the art in performance. For tests with unseen (but known) substrate a top-*k* approach is needed as only 4.3% of test cases predict the correct label as the nearest label. For top-10 that rises to 17%; for top-50 rises to 54%, and top-100 reaches 80%. TooTranslator demonstrates the potential of multimodal embedding alignment for open-world protein function inference.

## 1 Introduction

Many classification problems in protein sequence analysis suffer from a lack of data with the labels required for supervised learning. One way to address this is to use foundation Protein Language Models (pLMs), which use self-supervised learning on large amounts of unlabelled data, followed by transfer learning with a small labelled dataset. This approach follows the Closed-World Assumption (CWA), and can only predict membership in a class seen during training[BB15]. Open classification [SXL18], or novel class discovery [TLG^+^23], removes this assumption. It follows the Open-World Assumption (OWA) and classifies a test input as either belonging to a seen class, or as new.

Ideally, when you have a space of potential labels for a test input, even if most of them are not seen during training, you would like open classification to not only say the test input is new, but also to assign a label to it: a label unseen during training. This is the aim of zero-shot classification: the model is expected to predict new labels without prior training on them.

Transmembrane transport proteins are an essential group of proteins located in the membrane, transporting different substances intra- and inter-cellularly. They facilitate the selective passage of vital compounds ranging from small ions to macromolecules, through either active transport or diffusion. This biological process is an important part of nutrient acquisition, development, cellular homeostasis, cell metabolism, signaling and communication, and drug uptake and efflux. The substrates transported by these proteins can range from small ions such as sodium and potassium; sugars such as glucose; to larger compounds such as other proteins and messenger molecules [LBK^+^08]. Despite their biological and medical importance, experimental characterization of transporter specificity remains limited because membrane proteins are difficult to express and reconstitute *in vitro*, and substrate-binding assays are slow and labor-intensive [LXHNE17, Pol10]. As a result, only a small fraction of transporters in major databases such as UniProt-KB [The23] have experimentally verified substrates, creating a need for computational methods to infer these relationships.

The classification of transmembrane transport proteins aims to predict the class of substrates which a transporter moves across a membrane. The class could be a set of related chemical compounds, such as `inorganic cations`, or `amino acids`, or the class could be a single specific substrate such as `calcium ion or glutamine`. In the latter case, our space of labels is the universe of chemical compounds used within the cell, such as those in the `ChEBI` ontology for Chemical Entities of Biological Interest.

Existing computational models predict substrate classes using supervised learning on handcrafted protein features [MCZ14, AAB20, AB22]. These approaches rely on predefined substrate categories and require balanced training data to perform reliably. Although effective for well-studied transporter families, they struggle to generalize to novel proteins, uncommon substrates, or rare functional classes. The need to aggregate chemically diverse substrates into broad categories (e.g. cations or sugars) further limits their biological interpretability, since many molecules within a class differ substantially in physicochemical and pharmacological properties. The main limitation remains annotated data scarcity, as most specific substrates have very few known transporter examples.

Recent advances in pre-trained language models for proteins and chemical compounds enable representation learning directly from large unlabeled corpora [WLZ^+^25, RMS^+^21, EHD^+^22, ASC^+^22]. In this study, we explore whether mapping between these representation spaces can help infer substrate associations. This idea is conceptually related to the cross-lingual embedding alignment introduced by Mikolov et al. [MLS13], where word vector spaces from different languages were shown to be mappable through linear transformations. We extend this notion to the biological domain and ask whether a similar mapping can be learned between protein and chemical embedding spaces.

We propose TooTranslator, a model that aligns embeddings from ProtBERT (protein sequences) [EHD^+^22], ChemBERTa (molecular SMILES) [ASC^+^22], and SciBERT (descriptions) [BLC19] within a shared latent space. We selected three BERT-based models to maintain architectural and pretraining consistency, reducing the complexity of cross-domain mapping. TooTranslator translates between protein and molecule representations, allowing biochemical compatibility to be inferred from spatial proximity.

This approach supports few-shot learning (from few training samples in each class) and potentially zero-shot learning (from no training sample for unseen classes) substrate prediction by reasoning through semantic (embedding) similarity. While the model currently shows limited generalization to unseen substrates, it provides an initial exploration of translating representations between proteins and small molecules and highlights key challenges in learning cross-domain biological semantics.

When classifying specific substrates of transporters [AB25], we constructed a dataset from UniProt that identified transport proteins annotated with 149 different specific substrates There were 96 substrates with 3 or more known transport proteins, and 53 substrates with 1 or 2 known proteins; and many transport proteins for which the substrate was not known.

This paper aims to evaluate whether training the model on a limited set of protein-substrate pairs can lead to improved representations for unseen classes, enabling an accurate zero-shot classification. Specifically, we transform the problem by replacing the class labels with their detailed descriptions (`ChEBI` definitions and SMILES strings), which are then vectorized using two BERT encoders. This shifts the task from a traditional classification problem to a regression problem, where the model learns to translate protein vectors into label vectors. By training the model on a restricted dataset and updating the network weights for seen classes, we aim to determine if it can generalize to unseen substrates. The hypothesis is that samples from unseen classes will be closer to their label vectors in the learned embedding space, even without direct training.

We study multi-modal learning where three language models, ProtBERT-BFD, SciBERT, and Chem-BERTa, for three modes, protein sequence, scientific text, and chemical compounds, are unified into a single embedding space to facilitate zero-shot learning. The dataset and code for this project are publicly available on GitHub at the link: https://github.com/simaataei/TooTranslator.

The rest of this paper is organized as follows: Section 2 presents background material. Section 3 describes the structure of TooTranslator. Section 4 describes the Materials and Methods used in this work. Section 5 presents the results of our experimental evaluation. Section 6 has a discussion of the results. Finally, Section 7 concludes the paper.

## 2 Background

Recent advancements in Natural Language Processing (NLP) have been significantly driven by the development of Large Language Models (LLMs) [MRS08]. Central to these models are transformer architectures [Wol20], which utilize self-attention mechanisms to effectively capture and represent the contextual relationships within textual data [VSP^+^17]. A notable example is BERT [DCLT19], which has served as a foundational model influencing numerous subsequent architectures.

BERT (Bidirectional Encoder Representations from Transformers) [DCLT19] is a transformer-based model designed for natural language understanding. It features an encoder architecture with 12 layers and utilizes multi-head self-attention mechanisms to capture contextual relationships in text. The base version of BERT has a hidden size of 768 and contains around 110 million parameters. BERT is pre-trained on large datasets, including Wikipedia and BooksCorpus, enabling it to learn rich bidirectional representations [DCLT19].

There are two conventional transfer learning approaches [TS10] to using BERT-based language models in downstream classification tasks. First, a pre-trained language model could be utilized as an encoder, producing vectorized encoding for the input proteins. These encodings are fed into a classifier layer learning to predict the targets according to the specified task. This method is also known as encoding construction using a *frozen* language model. On the other hand, *fine-tuning* a BERT language model includes updating the weights of the model during the process of training to adjust the encoding according to the downstream task, which is classification in this case.

Protein sequences in their primary structure are represented as chains of amino acids, similar to how sentences are composed of words in a language. By viewing the 20 amino acids as an alphabet, Protein Language Models (pLMs) leverage advances in Natural Language Processing (NLP) to capture and represent the underlying information in protein sequences [OBL21]. The introduction of transformer models [VSP^+^17] learns contextual relationships between amino acids, allowing protein language models to capture structural, functional, and evolutionary information, enabling tasks such as protein classification, structure prediction, and functional annotation.

In 2019, Elnaggar et al. [EHD^+^22] introduced six protein language models by training two auto-regressive models (Transformer-XL, XLNet) and four auto-encoder models (BERT, Albert, Electra, T5). They trained each model on UniRef and the Big Fantastic Database (BFD) on the Summit supercomputer using 5616 GPUs and TPU Pod up to 1024 cores. BFD [JEP^+^21] is created by clustering 2.5 billion protein sequences from UniProt, Metaclust, and the Soil Reference Catalog Marine Eukaryotic Reference Catalog. It represents one of the largest publicly available collections of protein families, consisting of 65,983,866 families covering 2,204,359,010 protein sequences.

The ProtBERT model they introduced is a BERT-based protein language model. The advantage of using ProtBERT’s encoding vectors as exclusive input for several downstream tasks is validated in this paper. Although not the best language model in this study, the model is widely used due to its accessibility, practical model size, and usability in fine-tuning tasks.

ProtBERT-BFD is used for our study. It is a BERT model trained on BFD using Masked Language Modeling (MLM) task only. It has more layers than the original BERT. The model has 30 layers, 16 attention heads, and 420M parameters. ProtBERT-BFD was first trained for 800k steps for sequences with a maximum length of 512, and then for another 200k steps for sequences with a maximum length of 2k, in order to first extract useful features from shorter sequences while using a larger batch size. The optimizer used is Lamb with a learning rate of 0.002, a weight decay of 0.01, a learning rate warm-up for 140k steps then followed by linear decay of the learning rate.

### 2.1 Inputs and Encodings

The inputs for TooTranslator are protein sequences given in `fasta` format; a short text description of a substrate taken from the `Definition` field of its `ChEBI` entry; and the SMILES text formula [Wei88] of the substrate taken from the `SMILES` field. For example, the entry `CHEBI:33551`, `organosulfonic acids` has definition *“Organic derivatives of sulfonic acid in which the sulfo group is linked directly to carbon”* and SMILES string *“OS([*^*\**^*])(=O)=O”*.

The vectorization of the inputs is done by three BERT-based models: ProtBERT-BFD [EHD^+^22]; SciBERT [BLC19]; and ChemBERTa [ASC^+^22].

ProtBERT-BFD is described above. SciBERT has 12 layers and 12 attention like the original BERT. It was trained on 1.14 M full-text scientific papers, including the biomedical domains, from Semantic Scholar. SciBERT used a custom “wordpiece vocabulary” created to better suit the scientific text corpus, making it more effective for domain-specific tasks than the general BERT model. ChemBERTa has 6 layers and 12 attention heads for encoding SMILES strings. It used MLM training that masked 15% of the SMILES string. It was trained on the PubChem dataset which contains over 111 M chemical compounds in SMILES format.

### 2.2 Imbalance

The assumption of a uniform distribution for samples in the classes of the dataset does not hold in many real-world datasets. Imbalance is often a problem. An *imbalanced dataset* has a non-uniform distribution of instances across different classes, resulting in one or more classes being significantly overrepresented (majority classes) while others are underrepresented (minority classes). The ratio of imbalance is defined with the ratio of the smallest to the largest class as *s* : *l*, with *s* samples in the smallest and *l* samples in the largest class [HG09]. This ratio helps categorize imbalanced datasets into three groups: *slight imbalanced, extreme imbalanced*, and *severe imbalanced* datasets, where the ratio is less than 1:4, between 1:4 to 1:100, and over 1:100 respectively [Kra16].

Class imbalance could occur in the datasets of both binary and multi-class classification problems. Studying multi-class imbalanced datasets is not a straightforward task due to the multiple relationships between classes in multi-class problems; for example, a class might be majority compared to one class, but balanced compared to another class in the dataset [SG12]. Most of the research on class imbalance is developed for the binary classification problems leaving multi-class imbalance a relatively under-studied subject. In addition, the majority of this research work is on *slight imbalance*, resulting in under-developed *severe imbalance* problems [ZXL^+^16].

*Long-tailed imbalance* is a specific type of class imbalance in which a multi-class dataset demonstrates a long-tailed distribution, characterized by a few majority classes with many samples and numerous minority classes with only a few samples [ZKH^+^23]. The task of specific substrate classification [AB25] is a multi-class classification problem, that exhibits severe imbalance and long-tailed imbalance.

### 2.3 Learning from imbalanced data

To solve the problem of class imbalance, three main approaches have been applied: data-level methods, algorithm-level methods, and hybrid methods [Kra16]. Data-level methods modify the collection of examples to balance distributions by resampling; introducing approaches that generate new objects for minority groups (oversampling); and that remove examples from majority groups (undersampling) [FGHC18, MRA20, LWZ08]. Algorithm-level methods directly modify existing learning algorithms to alleviate the bias toward the majority classes. The main approaches are cost-sensitive learning, contrastive learning, and ensemble learning [ZL10, MMB21, KWH15]. Hybrid methods combine the advantages of two previous methods: merging sampling and cost-sensitive learning techniques [WGC14].

The main concern with the severe class imbalance in small minority classes is the risk of poor representation and lack of clear structure in small classes. In such cases, the straightforward application of oversampling methods that rely on relations between minority objects could result in overfitting the new samples on the weakly represented distribution of current samples [KKP06]. Another obstacle to resampling methods is the overlapping representation of different classes in the dataset. The overlapping of samples from different classes results in the unreliability of the basic resampling techniques.

### 2.4 Low-shot Learning

*Few-shot learning* methods aim to solve a supervised task with only a small number of training samples for certain classes in the training set [BPPK24]. *Low-shot learning* is used as an umbrella term for a model to learn information from few, one, or zero samples from a specific class in the training set [WYKN20]. The motivation for developing low-shot learning models lies in the fact that these models have many useful applications in real-world problems. Many real-world problems suffer from scarce data samples due to data acquisition and manual labeling process drawbacks. Therefore, low-shot learning models that generalize from a few examples are a matter of interest [BMR^+^20, LEB08].

### 2.5 ProTranslator

ProTranslator [XW22] is a zero-shot classification model developed to solve the task of protein function prediction. This model relates proteins to functions based on the textual descriptions available for the functions. Redefining the problem as a regression task using metric learning [BHS15], the authors propose a model to translate the description word sequence of a function into the amino acid sequence of a protein. By transferring annotations from functions with similar textual descriptions, ProTranslator can annotate novel functions.

ProTranslator does the following:

▸ Encode a vector representation for Gene Ontology (GO) functions based on their textual descriptions, using PubMedBERT [GTC^+^21].
▸ Encode proteins based on sequence (with CNN), textual descriptions from GeneCards (with BERT model), and protein-protein interaction (PPI) networks (with GNN).
▸ Combine features by processing sequence, description, and network encodings through individual one-depth fully connected layers to ensure their dimensions are uniform. The vectors are concatenated into one protein vector, which is projected into a shared low-dimensional encoding space along with GO function vectors.
▸ Predict protein function by calculating similarities between the projected GO term encodings and protein encodings in the shared space.

It outperforms state-of-the-art methods especially in zero-shot predictions, demonstrating its capability to generalize to unseen functions. ProTranslator relies heavily on the availability of textual descriptions for new functions, and on high-quality PPI network data. These may be incomplete or unreliable.

### 2.6 Loss Functions

The goal of training is to minimize the calculated loss by updating the model’s parameters. Different loss functions are employed in the training process of the models in various scenarios: Mean Squared Error (MSE) loss, weighted Mean Squared Error loss, Mean Absolute Error (MAE) loss [Ber13], and Huber loss [Hub92].

#### Mean Squared Error and weighted Mean Squared Error

Mean Squared Error (MSE) or L2 loss calculates the average squared difference between the model’s predicted values and the actual values. For *n* number of samples, *y* as the actual value, and *ŷ* as the predicted value of each sample, the MSE loss is calculated as follows:

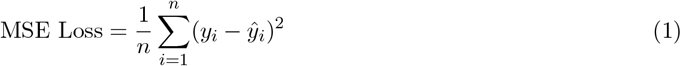

In many cases, different classes are valued differently during the training process. For an imbalanced dataset, classes with a lower number of samples are given greater priority compared to the majority classes. To address this, a weight 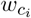 is applied to each class *c*_*i*_ in the MSE loss to account for the varying significance, as shown in Equation 2 [Ber13]:

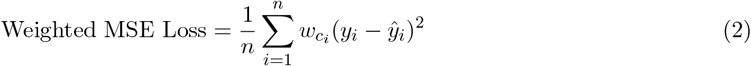

#### Mean Absolute Error

Mean Absolute Error (MAE) or L1 loss calculates the average of the absolute difference between the model’s predicted values and the actual values. In contrast to MSE loss, which penalizes larger errors more heavily and is sensitive to outliers, MAE loss does not square the errors, treating all deviations equally. This characteristic makes MAE more robust to outliers in the data. Therefore, MSE is preferred when the goal is to emphasize larger errors and when outliers are a significant aspect of the problem. On the other hand, MAE is useful when dealing with data containing outliers or when a more consistent, robust error measure is desired. For *n* number of samples, *y* as the actual value, and *ŷ* as the predicted value of each sample, the MAE loss is calculated as follows:

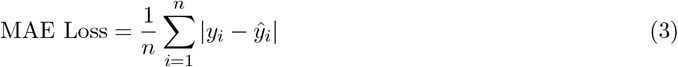

#### Huber Loss

The Huber loss function is introduced to address challenges in the robust estimation of location parameters, making the estimator less sensitive to outliers. This function has combines Mean Squared Error (MSE) and Mean Absolute Error (MAE) loss functions, which is useful where the sample mean is excessively influenced by a few extreme values in long-tailed distributions. Defining *y* as the actual value, *ŷ* is the predicted value for a sample, and *δ* as a constant threshold, the Huber loss function is quadratic for small value differences and linear for larger value differences [Hub92]:

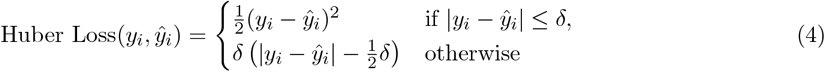

## 3 TooTranslator Structure

The TooTranslator model “translates” the primary structure of transmembrane transport proteins to their carried substrate. This model modifies the protein sequence encoder weights based on seen *<*protein sequence, label*>* pairs aiming to produce reliable encoding vectors for the unseen classes of proteins.

The workflow consists of two stages: (1) **Vectorization and Projection**, where protein and substrate embeddings are generated and projected into the shared space; (2) **Substrate Retrieval**, where candidate substrates are ranked according to their distance to the projected protein embedding.

TooTranslator (Figure 1) employs three language models for vectorization: ProtBERT-BFD [EHD^+^22] for protein encoding; SciBERT [BLC19] for encoding substrate descriptions; and ChemBERTa [ASC^+^22] for encoding the substrate’s SMILES string.

**Figure 1.**
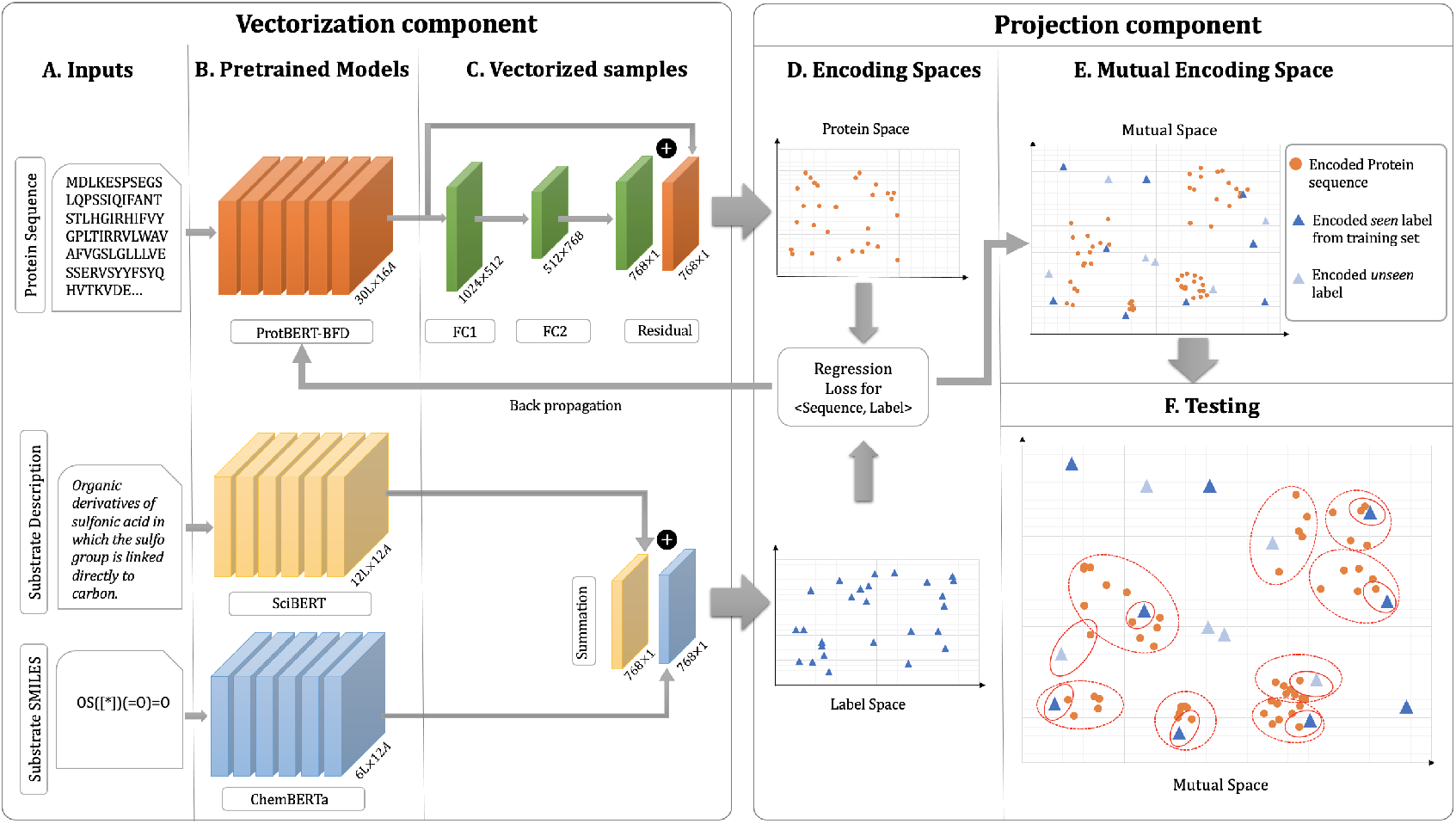
TooTranslator model with two components: vectorization and projection. The figure presents the training and testing process for the TooTranslator model. In the training phase, the samples are a pair of *<*protein sequence, label*>*. Each label includes a substrate definition and a SMILES string (A). Protein sequences are encoded using ProtBERT-BFD pLM. The label definitions and SMILES strings are encoded using SciBERT and ChemBERTa-2 respectively (B). L and A show the number of layers and attention heads in each model and the size of the layers and output vectors is presented under each. Two fully connected layers and one residual layer are applied to the protein vectors resulting in a vector with the same size as labels. The label definition and SMILES vectors are summed element-wise into one vector representing the label (C). Vectorizing all of the samples in the training set results in two encoding spaces: protein space and label space (D). The protein space is projected into the label space calculating a regression loss for sequence, and label vector pairs; back-propagating the loss into the ProtBERT-BFD pLM. The mutual space includes both protein and label vectors (E). This mutual space also includes unseen labels from `ChEBI` (light blue triangles) vectorized without any proteins associated with them. In the testing phase, each test sample is a protein sequence in the mutual space. The nearest label vector to the protein in this space is selected as the prediction (F).

The projection process involves mapping the protein-encoding space to the label-encoding space, to minimize the distance between the protein-encoding vector and its corresponding substrate-encoding vector. To identify the best architecture for this mapping task, various combinations of fully connected layers, residual networks, and multi-head attention layers were explored. The optimal network architecture consists of two fully connected layers followed by a residual layer [HZRS16] (Figure 1). The first fully connected layer (FC1) maps the 1024-dimensional output vectors from ProtBERT-BFD to a 512-dimensional space. The second fully connected layer (FC2) maps this vector from 512 to 768 dimensions. In the residual layer, the initial output of the ProtBERT-BFD model (truncated to 768 dimensions) is added to the final output of the second fully connected layer (FC2).

Classification is performed by substrate retrieval. A test protein is encoded via ProtBERT-BFD and projected into the shared latent space Distances between the projected protein vector and all fixed substrate embeddings are computed using the same metric as in training. The nearest substrate vector is selected as the predicted candidate. This approach allows predictions for unseen substrate classes, since each substrate embedding is independently vectorized.

## 4 Materials and Methods

### 4.1 Datasets

Our previous study [AB25] constructed a dataset from UniProt using the Gene Ontology (GO) Molecular Function (MF) terms rooted at *“GO:0022857”* (transmembrane transporter activity) and their corresponding `ChEBI` terms. We selected leaf terms so the substrates were specific substrates rather than substrate collections to relate proteins and substrates. Duplicates were removed. A threshold of 3 samples per substrate was set for the dataset UniProt-SPEC-100 used in that study. It had 96 specific substrates and 4,455 proteins. This excluded 70 proteins and 53 specific substrates where the threshold was not met.

For the previous study, as the smallest class has size three, we split the dataset on 34% to 66% test-train ratio. This ratio secures one sequence from the smallest class for the stratified test-train split. That gave a hold-out test set with 1,485 proteins and a training set with 2,970 proteins.

For this TooTranslator study we use three datasets:

**Dataset 1 (DS1)** is the above training set with 2,970 proteins and 96 seen substrate classes, which is used for training and cross-validation;

**Test Set 1 (TS1)** is the above hold-out test set of seen classes with 1,485 proteins and 96 substrate classes; and

**Test Set 2 (TS2)** is the excluded set of 70 proteins and 53 unseen specific substrates.

### 4.2 Training

Training TooTranslator occurs during the projection process, where the protein space is mapped to the label space. In this phase, for each input sequence, the protein-encoding vector is extracted from the ProtBERT-BFD language model, followed by two fully connected layers and a residual layer (Figure 1). Simultaneously, the corresponding label description is encoded by SciBERT, and the SMILES string is encoded by ChemBERTa. The final label encoding vector is the summation of the two vectors.

The goal of training is to minimize the loss between the protein-encoding vector and the label-encoding vector (Figure 1D). The loss is calculated and back-propagated through the ProtBERT-BFD network. Updating the weights of the ProtBERT-BFD model improves the protein encodings, bringing the protein vectors in the substrate space closer to their corresponding substrate labels (Figure 1E). To identify the optimal loss function for this process, four separate models are implemented, each with a different loss function: mean squared error (MSE loss), weighted mean squared error (WMSE loss), mean absolute error (MAE loss), and Huber loss.

The models are developed in `Python`, version 3.9.1, using `PyTorch` and `Transformers` [PGM^+^19, WDS^+^20]. For all four models, the training parameters are set as follows: 100 epochs of training with batch size 1, AdamW optimizer, and learning rate equal to 5e-6 [KB15]. To find the best hyperparameters for the TooTranslator model trained with Huber loss, the following search space is explored: number of epochs from 1 to 200; batch size: 1, 2, 10, and 40; and learning rate between 1e-6 to 1e-2. All models are trained and tested on Concordia University’s High-Performance Computing (HPC) Facility (`speed`) using one NVIDIA Tesla P6 GPU, with 16 GB of memory.

### 4.3 Testing

In classification, a protein sequence is presented to TooTranslator as a test sample. The trained ProtBERT-BFD encodes this sample into a vector, which is then passed through a network structure with fully connected and residual layers, resulting in a mutual space with 768 dimensions. In this space, the pairwise distance between the protein vector and existing label vectors is calculated, and the nearest label vector to the protein vector is selected as the prediction for the test sample.

This differs slightly from K-Nearest Neighbors (KNN), where the labels of the K nearest protein samples are retrieved as the prediction. In TooTranslator, classification can include label vectors from unseen classes, as the label space includes all substrates in ChEBI. The unseen classes could be the predictions of novel proteins. In practice, scientists require top-k prediction results in order to design and prioritise their wet lab experiments.

This model is tested on two separate test sets. **TS1** is the hold-out set derived from the original train/test split of the dataset, with a proportion of 2:1. It contains only seen classes from the training set. **TS2** evaluates the model’s performance on unseen classes. It contains classes that were excluded from the original dataset for not meeting the sample size threshold of three.

### 4.4 Evaluation Metrics

Performance evaluation of the classification models considers five metrics:

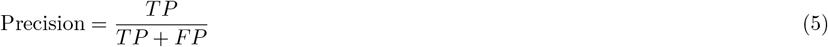

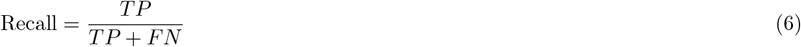

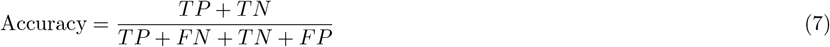

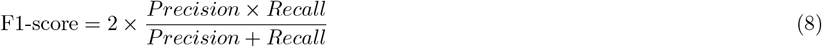

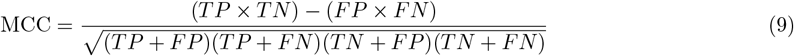

where *TP* is the number of true positives, *TN* is the number of true negatives, *FP* is the number of false positives, and *FN* is the number of false negatives. For imbalanced data, the Matthews Correlation Coefficient (MCC) is arguably the best single assessment metric [Din11, WP03, BDA13]. The overall performance across all classes is the micro-average of the individual results [MRS08] and we used the multi-class version of MCC [Gor04].

## 5 Results

Results are presented for cross-validation (with seen classes) on the training set **DS-1**, the hold-out test set **TS1** of seen classes, and the hold-out test set **TS2** of unseen classes. Five metrics, accuracy, precision, recall, F1-score, and MCC, are reported. F1-score and MCC are the main metrics for evaluation of the loss functions, but precision and recall can be important in other scenarios.

### Results on seen classes

Table 1 compares the overall F1-score and MCC metrics for the four models using Huber loss, MSE loss, wMSE loss, and MAE loss functions. It also shows the number of seen classes that could not be predicted by these models. Table 2 presents the 3-fold cross-validation results for each model and permits a one-way ANOVA test. The last row shows the ANOVA p-value for each metric. They are all larger than 0.05, indicating that there is no statistically significant difference in model performance. However, the TooTranslator trained with the Huber loss function is selected due to its low number of unpredicted classes. Table 4 presents the five metrics across **TS1** for each seen class.

**Table 1:** Results of TooTranslator trained with four loss functions on TS1.

| Model name | Loss function | F1-score | MCC | Unpredicted |
| --- | --- | --- | --- | --- |
| TooTranslator-MSE | Mean Square Error | 0.740 | 0.908 | 7 |
| TooTranslator-wMSE | Weighted Mean Square Error | 0.748 | 0.918 | **3** |
| TooTranslator-MAE | Mean Absolute Error | 0.762 | **0.928** | 7 |
| TooTranslator-Huber | Huber | **0.790** | 0.926 | **3** |

**Table 2:**
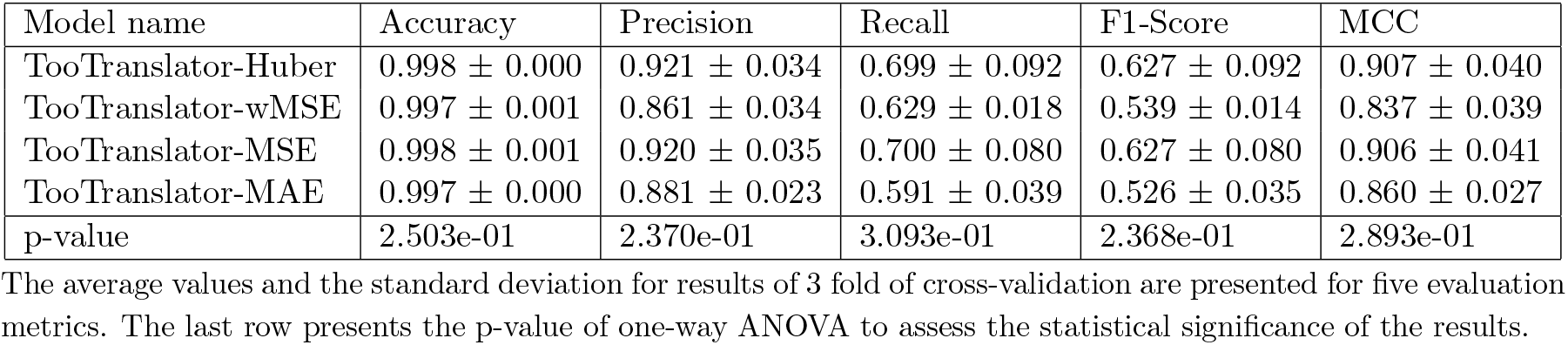
3-Fold Cross-validation results for TooTranslator with four loss functions.

| Model name | Accuracy | Precision | Recall | F1-Score | MCC |
| --- | --- | --- | --- | --- | --- |
| TooTranslator-Huber | $0.998 \pm 0.000$ | $0.921 \pm 0.034$ | $0.699 \pm 0.092$ | $0.627 \pm 0.092$ | $0.907 \pm 0.040$ |
| TooTranslator-wMSE | $0.997 \pm 0.001$ | $0.861 \pm 0.034$ | $0.629 \pm 0.018$ | $0.539 \pm 0.014$ | $0.837 \pm 0.039$ |
| TooTranslator-MSE | $0.998 \pm 0.001$ | $0.920 \pm 0.035$ | $0.700 \pm 0.080$ | $0.627 \pm 0.080$ | $0.906 \pm 0.041$ |
| TooTranslator-MAE | $0.997 \pm 0.000$ | $0.881 \pm 0.023$ | $0.591 \pm 0.039$ | $0.526 \pm 0.035$ | $0.860 \pm 0.027$ |
| p-value | 2.503e-01 | 2.370e-01 | 3.093e-01 | 2.368e-01 | 2.893e-01 |

The average values and the standard deviation for results of 3 fold of cross-validation are presented for five evaluation metrics. The last row presents the p-value of one-way ANOVA to assess the statistical significance of the results.

**Table 3:**
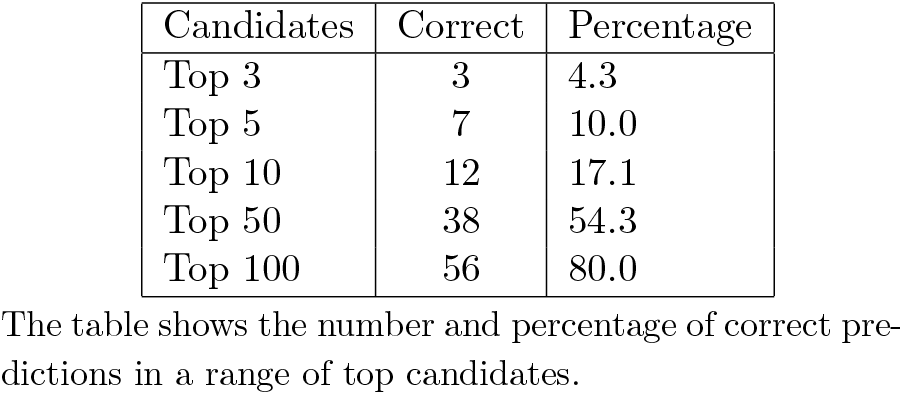
Top-k results for TS2.

| Candidates | Correct | Percentage |
| --- | --- | --- |
| Top 3 | 3 | 4.3 |
| Top 5 | 7 | 10.0 |
| Top 10 | 12 | 17.1 |
| Top 50 | 38 | 54.3 |
| Top 100 | 56 | 80.0 |

The table shows the number and percentage of correct predictions in a range of top candidates.

**Table 4:**
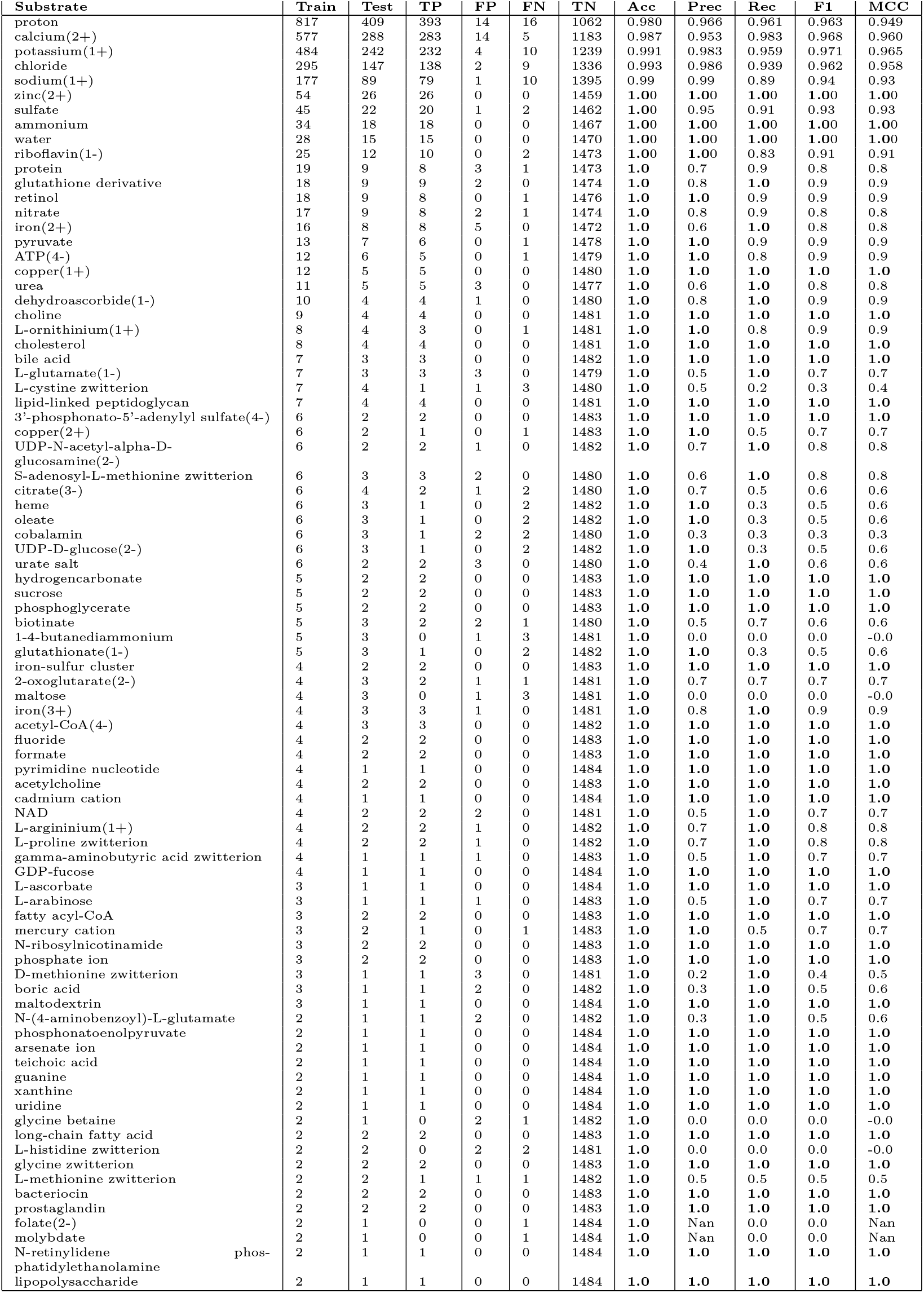
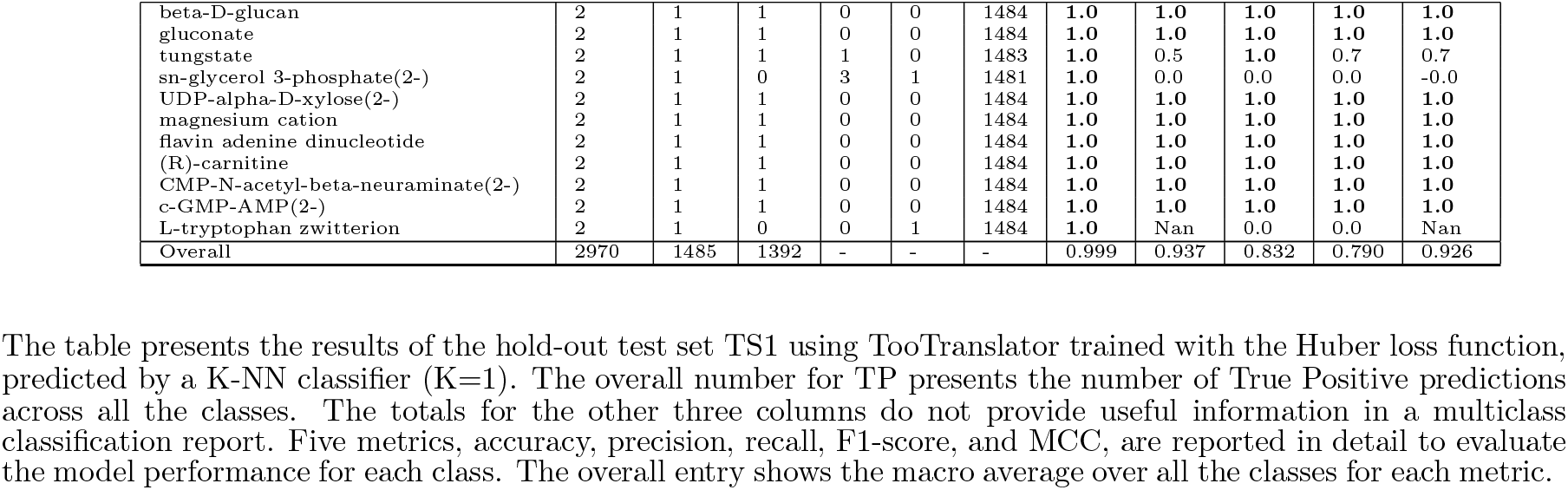
TooTranslator with Huber loss function results for TS1.

### Results on unseen classes

For the test samples in **TS2**, although the substrates were not seen during training, we do know the actual substrate. Table 5 presents the TooTranslator predictions for each test sample along with the correct label. It also shows the distances of the protein sequence projection from the true labels and the prediction. The last column indicates that only three test samples have the predicted label matching the actual label.

**Table 5:**
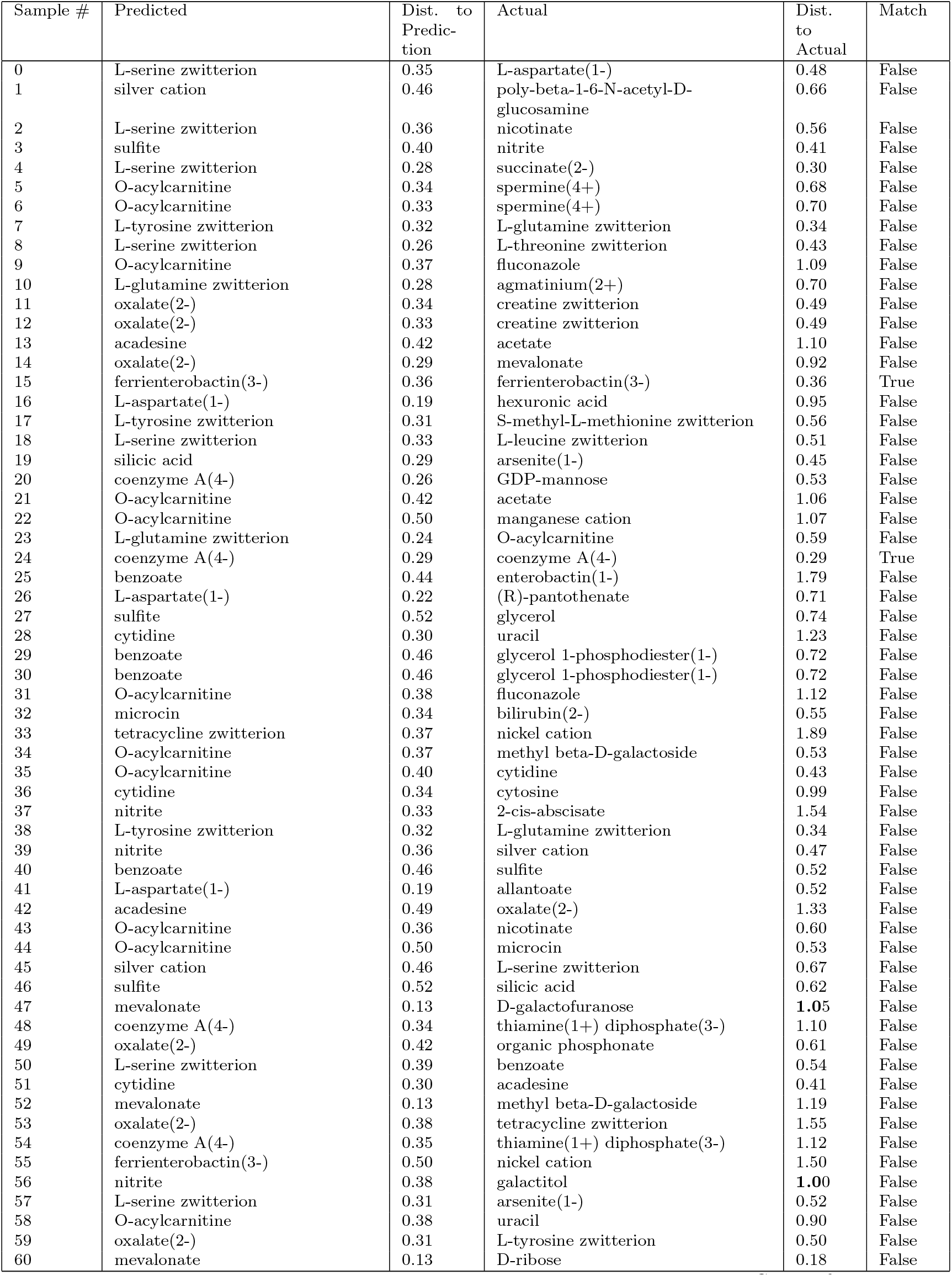
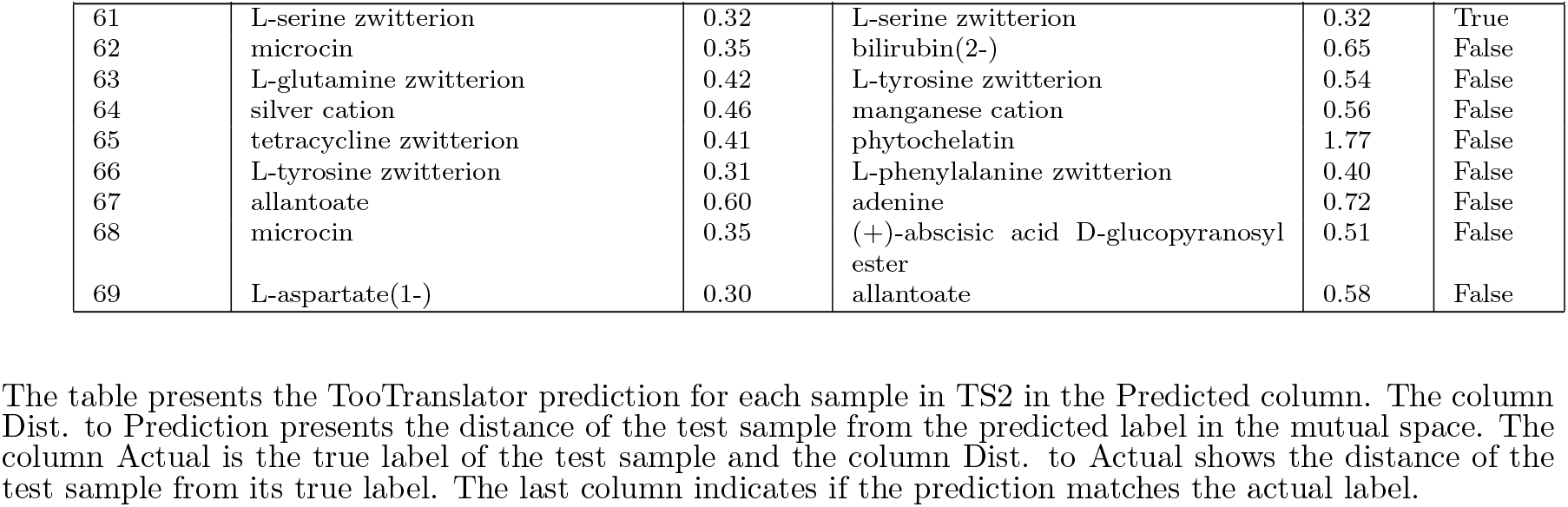
Results and Distances of TooTranslator with Huber loss for TS2.

Table 6 presents the labels of incorrect predictions for unseen classes in **TS2**, ordered by frequency of occurrence. There are 33 labels, 2 are unseen classes, and the remaining 31 are seen during training.

**Table 6:** TooTranslator predicted labels for TS2.

| # | Label | Freq. | Seen |
| --- | --- | --- | --- |
| 0 | (R)-carnitine | 6 | seen |
| 1 | L-serine zwitterion | 5 | unseen |
| 2 | D-methionine zwitterion | 4 | seen |
| 3 | pyruvate | 4 | seen |
| 4 | proton | 3 | seen |
| 5 | L-arabinose | 3 | seen |
| 6 | folate(2-) | 3 | seen |
| 7 | L-argininium(1+) | 3 | seen |
| 8 | ATP(4-) | 3 | seen |
| 9 | L-cystine zwitterion | 2 | seen |
| 10 | sn-glycerol 3-phosphate(2-) | 2 | seen |
| 11 | biotinate | 2 | seen |
| 12 | 1-4-butanediammonium | 2 | seen |
| 13 | urate salt | 2 | seen |
| 14 | cobalamin | 2 | seen |
| 15 | water | 2 | seen |
| 16 | L-ornithinium(1+) | 2 | seen |
| 17 | tungstate | 2 | seen |
| 18 | L-glutamate(1-) | 2 | seen |
| 19 | gamma-aminobutyric acid zwitterion | 2 | seen |
| 20 | L-proline zwitterion | 2 | seen |
| 21 | nitrate | 1 | seen |
| 22 | copper(1+) | 1 | seen |
| 23 | CMP-N-acetyl-beta-neuraminate(2-) | 1 | seen |
| 24 | calcium(2+) | 1 | seen |
| 25 | L-methionine zwitterion | 1 | seen |
| 26 | urea | 1 | seen |
| 27 | glutathione derivative | 1 | seen |
| 28 | iron(2+) | 1 | seen |
| 29 | acetyl-CoA(4-) | 1 | seen |
| 30 | guanine | 1 | seen |
| 31 | UDP-D-glucose(2-) | 1 | seen |
| 32 | silicic acid | 1 | unseen |

The table presents the labels of the incorrect predictions of samples in TS2 with the frequency of each label in incorrect predictions. The last column indicates whether the incorrect label was seen during training or not.

## 6 Discussion

We analyse how well the TooTranslator model predicts specific substrates for seen and unseen substrates. The model successfully maps the protein-encoding space to the label-encoding space for seen classes, and improves protein-label distances for unseen classes.

### No difference between the four loss functions

The one-way ANOVA analysis of the performance of the four models in Table 2 shows a p-value of 0.3, which is larger than 0.05 required to show statistical significance of the performance differences.

### On seen classes TooTranslator is comparable to state-of-the-art *TooT-SS*

Table 1 shows that TooTranslator with Huber loss function on the hold-out test set **TS1** of seen classes has an F1-score of 0.790 and MCC of 0.926, and is unable to predict 3 of the 96 classes. This compares well with the performance of *TooT-SS* [AB25]: F1-score 0.923, MCC 0.908, and 13 of 96 classes not predicted.

### For unseen classes, the correct label is predicted only 4.3% of the time

Table 5 shows that only 3 (4.3%) of the 70 test samples of **TS2** have the correct label as the nearest prediction. Hence, consideration of top-k performance is done.

### Correct label is in top-100 80% of the time

Deeper analysis looks at distance between predicted label and actual label for test samples in **TS2** and the rank of the correct label amongst those predictions. Table 3 looks at the top 3, 5, 10, 50, and 100 candidates of each test sample, and shows the number and percentage of times the correct label occurs amongst the top *n* candidates. So 12 test samples from **TS2** have their correct label among the top 10 substrate candidates, and 56 have their correct label within the top 100 candidates, which is 80% of the test samples.. Figure 2 presents the location of the correct label as a histogram in rank intervals from 1 to 150, with 10-step intervals. The largest number of correct predictions (12 test samples) occurs within the top 10 candidates. An additional 7 test samples have their correct predictions within ranks 11–20, while 8 test samples have correct predictions in the 21–30 range. The furthest correct prediction is in rank 131–140.

**Figure 2.**
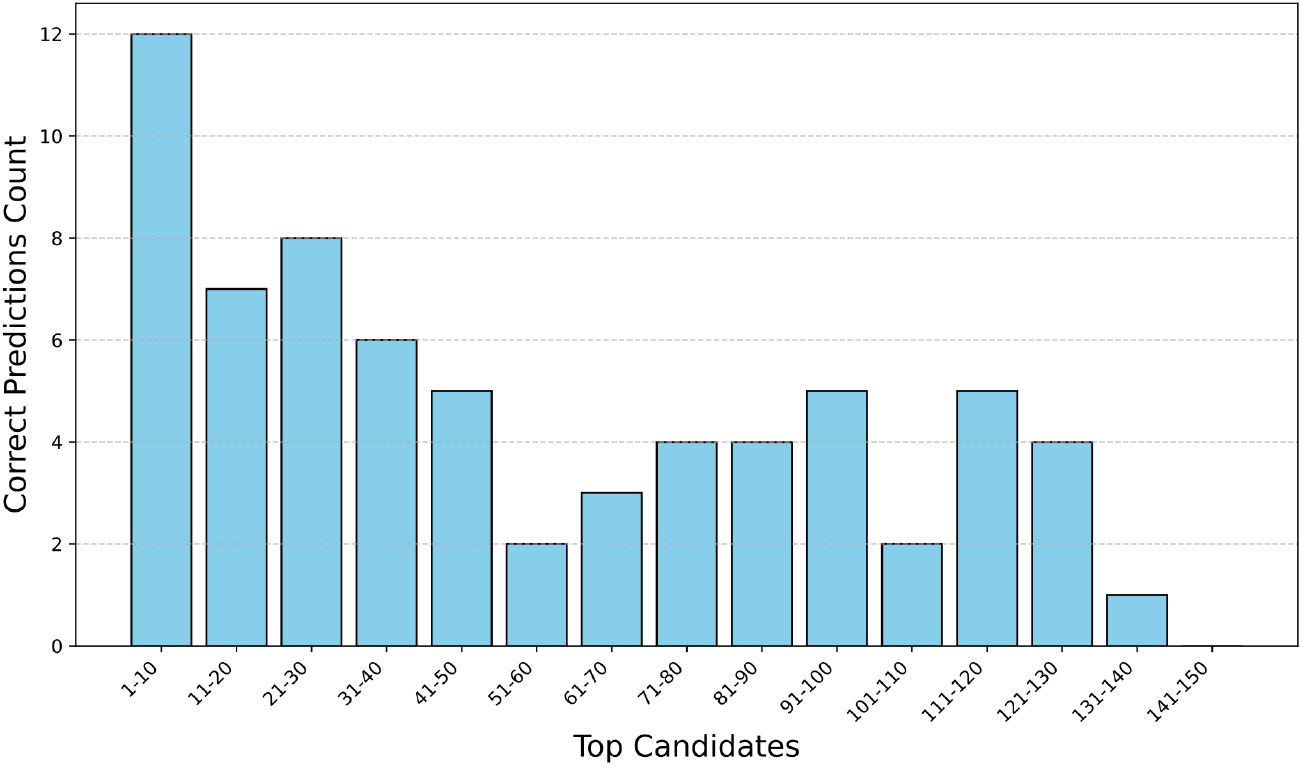
Range distribution of correct prediction for samples in TS2. The figure shows the number of correct predictions in top candidates for samples in TS2. The histogram ranges from 1 to 150 top candidates with 10-step intervals.

### Predicting 50% of unseen classes correctly requires top-50 results

Table 3 shows that the top-50 results include the correct label 54% of the time.

### Training brings both seen and unseen classes closer to the projected protein

We analyse whether training does bring projected protein label closer to its correct label. It is expected for proteins with seen labels as they are processed during training, but what of the unseen labels?

Figure 3 illustrates the representations of test samples from the seen classes during the training process (TS1). The objective of the study is to evaluate whether the training process updates the model weights to produce reliable representations of protein sequences from unseen classes (TS2), positioning them close to their corresponding label vectors.

**Figure 3.**
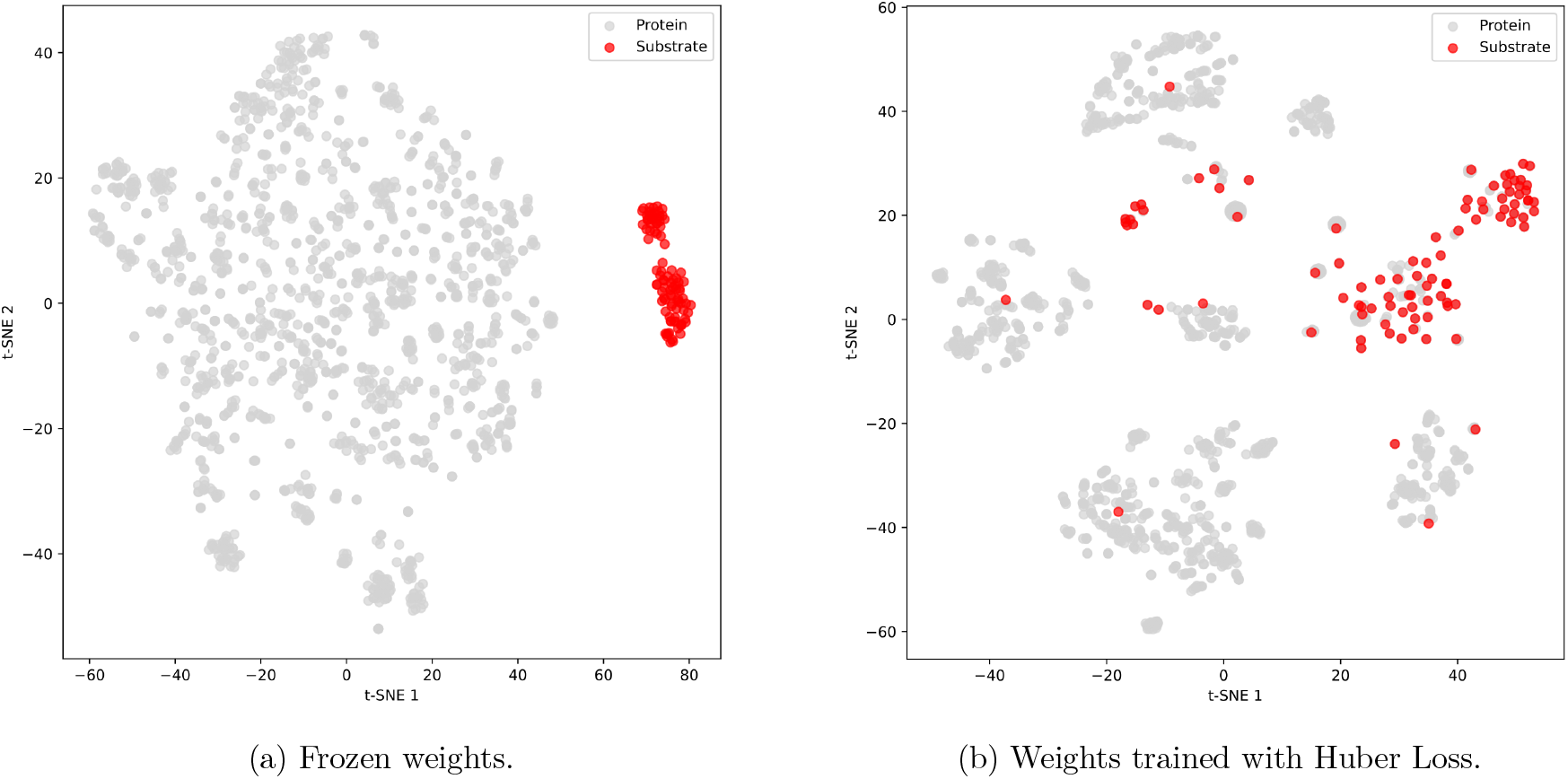
t-SNE plot for protein and label representations. The figure presents the 2-D t-SNE plot of protein representations (gray) and label representations (red) before and after training in TooTranslator with Huber loss for samples in TS1.

Figure 4 presents the distance distribution of samples in both TS1 and TS2 before and after training. It shows the distance of unseen classes from TS2 has decreased even though these classes are not included in the training process.

**Figure 4.**
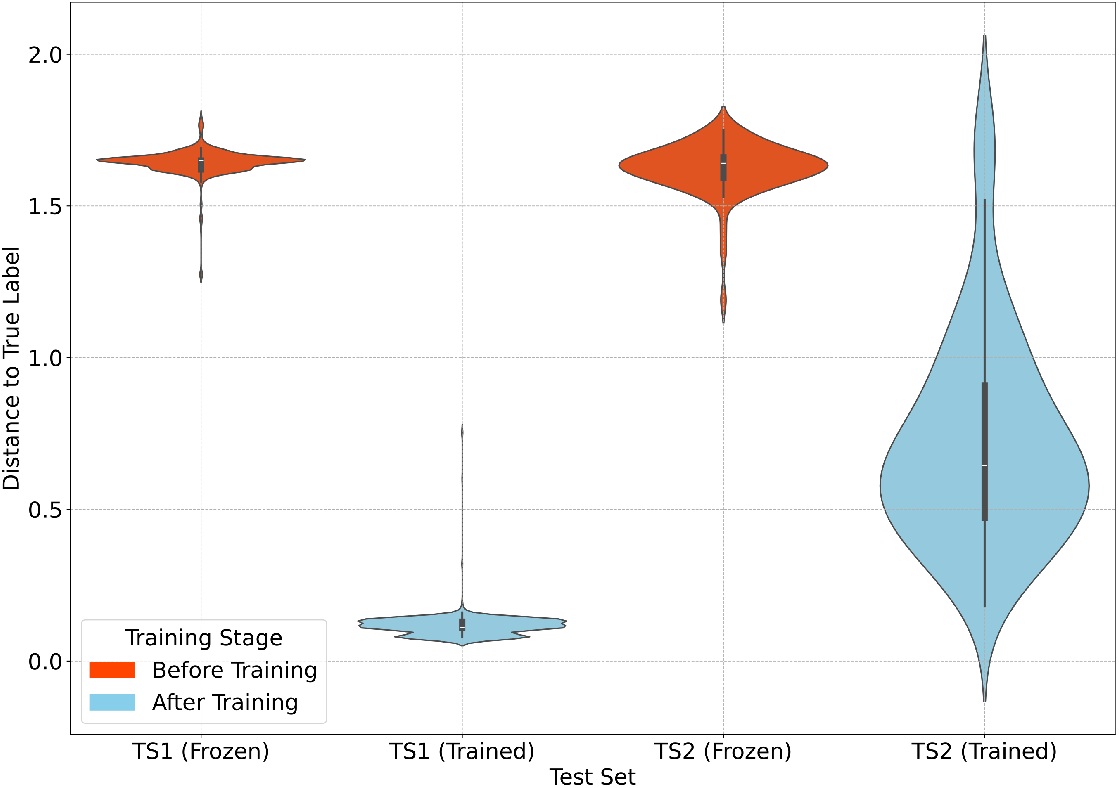
Distribution of distances to the true label before and after training. TooTranslator is tested with two test sets of TS1 and TS2. The figure shows the distribution of each test sample from its true label before (orange plot) and after (blue plot) training for TS1 (left) and TS2 (right).

## 7 Conclusion

This study addresses the challenge of identifying inorganic ion transport proteins and predicting their specific substrates using open-world classification to allow novel class discovery. The TooTranslator model is implemented to predict the specific substrate for an input protein, trained on the primary structure of the protein, and the chemical substrate’s `ChEBI` definition and SMILES string.

By redefining the classification problem as a regression task, TooTranslator aims to leverage the zero-shot learning process to predict groups of both seen and unseen classes. On MCC and the number of unpredicted classes, this model outperforms the state-of-the-art classification model for substrate specificity on seen classes [AB25]. However, the model’s top-10 performance on unseen classes predicts only 12 of 70 samples (17%) from **TS2** correctly.

The model successfully maps the protein-encoding space to the label-encoding space for seen classes. Two fully connected layers and one residual layer implement the mapping. A Huber loss function and the AdamW optimizer train the network. Loss back-propagation to the protein encoder produces accurate representations for samples in seen classes (Table 1, Table 4) and improves protein-label distances for unseen classes (Figure 4).

Although the model does not fully resolve the unseen class substrate prediction in the first substrate candidate, the top-100 correctly predicts 80% of unseen test samples. Improvements may come from refining the projection process to better map protein sequences and substrate encoding vectors, experimenting with alternative neural architectures such as graph-based networks or transformer variations, and incorporating multi-task learning to simultaneously learn substrate-specific and sequence-specific information. A key direction for future work is adjusting the training strategy to enable gradient back-propagation to both the label and protein encoders, which would require greater computational resources..

Since this is the first attempt in the field to predict specific novel substrates for new proteins, no existing models serve as direct baselines. While it does not fully solve the novel substrate identification problem, Figure 4 shows that training reduces the distance between unseen protein sequences and their true labels, despite these classes being absent during training. This highlights the model’s potential to generalize from known to novel substrate classes.

## Acknowledgment

This work was funded in part by Genome Canada, Genome Quebec, and Natural Sciences and Engineering Research Council of Canada (NSERC).

## Notes

https://github.com/simaataei/TooTranslator

### Competing Interest Statement

The authors have declared no competing interest.

